# Presynaptic PMCA interacts with Dlg, regulating in vivo synaptic calcium in *Drosophila* larval NMJ

**DOI:** 10.64898/2026.10.28.762485

**Authors:** Andrés Köhler-Solís, Estefanía López, Eduardo Quiroz, Oliver Kobler, Andrés González-Gutiérrez, Ulrich Thomas, Jimena Sierralta

## Abstract

Precise regulation of presynaptic Ca²⁺ is essential for neurotransmitter release and synaptic function. At the *Drosophila* larval neuromuscular junction (NMJ), the plasma membrane Ca²⁺-ATPase (PMCA) mediates presynaptic Ca²⁺ extrusion and is enriched in peri-active-zone membrane domains. Here, we investigated how PMCA distribution is regulated and contributes to synaptic function. Co-immunoprecipitation and super-resolution STED microscopy revealed that PMCA interacts and co-localizes with the MAGUK scaffold protein Discs large (Dlg) in discrete presynaptic membrane clusters. Neuronal depletion of the DlgS97 isoform disrupted PMCA clustering without substantially altering its abundance, redistributing the pump into smaller membrane puncta. Reporter assays further showed that PDZ-binding motifs in the PMCA C-terminus mediate its interaction with Dlg and are required for normal cluster formation. Electrophysiological analyses revealed that reducing either PMCA or Dlg increased spontaneous neurotransmitter release and enhanced asynchronous release following high-frequency stimulation. PMCA depletion also prolonged postsynaptic current decay during repetitive stimulation, implicating PMCA in the rapid regulation of presynaptic Ca²⁺ signals.

Together, these findings identify Dlg as a key organizer of presynaptic PMCA and establish a link between synaptic scaffold organization and Ca²⁺ homeostasis in vivo. Our results demonstrate that PMCA contributes to maintaining resting presynaptic Ca²⁺ levels, restoring Ca²⁺ homeostasis after repetitive stimulation, and regulating Ca²⁺ dynamics following a single stimulus. These findings highlight the importance of the nanoscale organization of Ca²⁺ extrusion machinery in controlling neurotransmitter release at the *Drosophila* NMJ.

## Introduction

Fast synaptic transmission depends on precise spatial and temporal regulation of intracellular Ca²⁺([1]). At chemical synapses, membrane depolarization triggers Ca²⁺ influx through voltage-gated calcium channels (VGCCs) concentrated at active zones, generating localized calcium nanodomains that initiate synaptic vesicle fusion and neurotransmitter release [2], [3]. In addition to these highly localized signals, residual and basal calcium levels regulate multiple aspects of synaptic function, including spontaneous release, vesicle priming, short-term plasticity, and asynchronous neurotransmitter release [4], [5], [6], [7], [8]. Consequently, presynaptic terminals require efficient mechanisms to restrict the spread and duration of calcium signals outside active zones.

Presynaptic calcium homeostasis is maintained through the combined action of calcium-binding proteins, intracellular sequestration systems, and plasma membrane calcium extrusion mechanisms. Among the latter, plasma membrane Ca²⁺ ATPases (PMCAs) and Na⁺/Ca²⁺ exchangers are major pathways for the removal of calcium from the cytoplasm to the extracellular compartment [9] [10] [11], [12]. Although both systems contribute to calcium extrusion, their relative importance varies considerably among neuronal types and synaptic compartments. At the *Drosophila* larval neuromuscular junction (NMJ), PMCA has been identified as the principal mechanism responsible for bulk calcium clearance following neuronal activity [13]. Recent work has further demonstrated that PMCA localizes predominantly within peri-active-zone membrane domains [14], [15],[16] where it limits the spread of calcium signals generated outside release sites and contributes to the functional separation of exocytotic and endocytotic calcium pathways [16].

Despite the recognized importance of PMCA in presynaptic calcium regulation, little is known about the molecular mechanisms that control its localization and organization within synaptic membranes. In several vertebrate cell types, PMCA isoforms interact with members of the membrane-associated guanylate kinase (MAGUK) family, including PSD-95/Dlg proteins, via C-terminal PDZ-binding motifs [17]. These interactions promote PMCA clustering and regulate its subcellular localization. PSD-95-dependent localization of PMCA in photoreceptor synapses [18], has been implicated in vitro in the control of calcium homeostasis, suggesting that scaffold-dependent organization of calcium extrusion machinery may represent a conserved mechanism for the regulation of synaptic calcium signalling.

The *Drosophila* MAGUK protein Discs large (Dlg) is a highly conserved scaffold protein known to control synaptic architecture and the localization of multiple membrane-associated proteins. At the larval NMJ, Dlg plays a role in the organization of presynaptic calcium channels and is required for normal synaptic transmission [19]. Moreover, we observed slower calcium recovery following depolarization in *dlg* mutant terminals [19], suggesting that Dlg may influence calcium homeostasis through mechanisms beyond its established effects on calcium influx. Dlg is expressed in two main splice variants, DlgA and DlgS97; DlgS97 is mainly neuronal, and DlgA is ubiquitous but far less expressed in neurons [21].

Here, we investigated the in vivo role of PMCA in regulating calcium-dependent synaptic function at the *Drosophila* larval NMJ and whether Dlg organizes presynaptic PMCA to support the formation of calcium nanodomains. Using biochemical approaches, super-resolution microscopy, genetic manipulation, and electrophysiology, we demonstrate that Dlg associates with PMCA in vivo and promotes its assembly into peri-active-zone membrane clusters. Disruption of this organization alters spontaneous and activity-dependent neurotransmitter release, supporting a role for Dlg-dependent PMCA clustering in presynaptic calcium homeostasis. Furthermore, we demonstrate that PMCA contributes to rapid calcium recovery following a single stimulus, highlighting its role in the precise regulation of calcium dynamics during neurotransmitter release.

## RESULTS

### Dlg and PMCA form a complex and co-localize at presynaptic terminals

To determine whether Dlg associates with PMCA in vivo, we performed co-immunoprecipitation experiments using adult head extracts from flies expressing endogenous genomically Venus-tagged PMCA (hereupon PMCA-YFP; [20]). Dlg was readily detected in PMCA-YFP immunoprecipitates but not in control preparations, indicating that both proteins are components of the same molecular complex (Fig. 1A).

**Figure 1:**
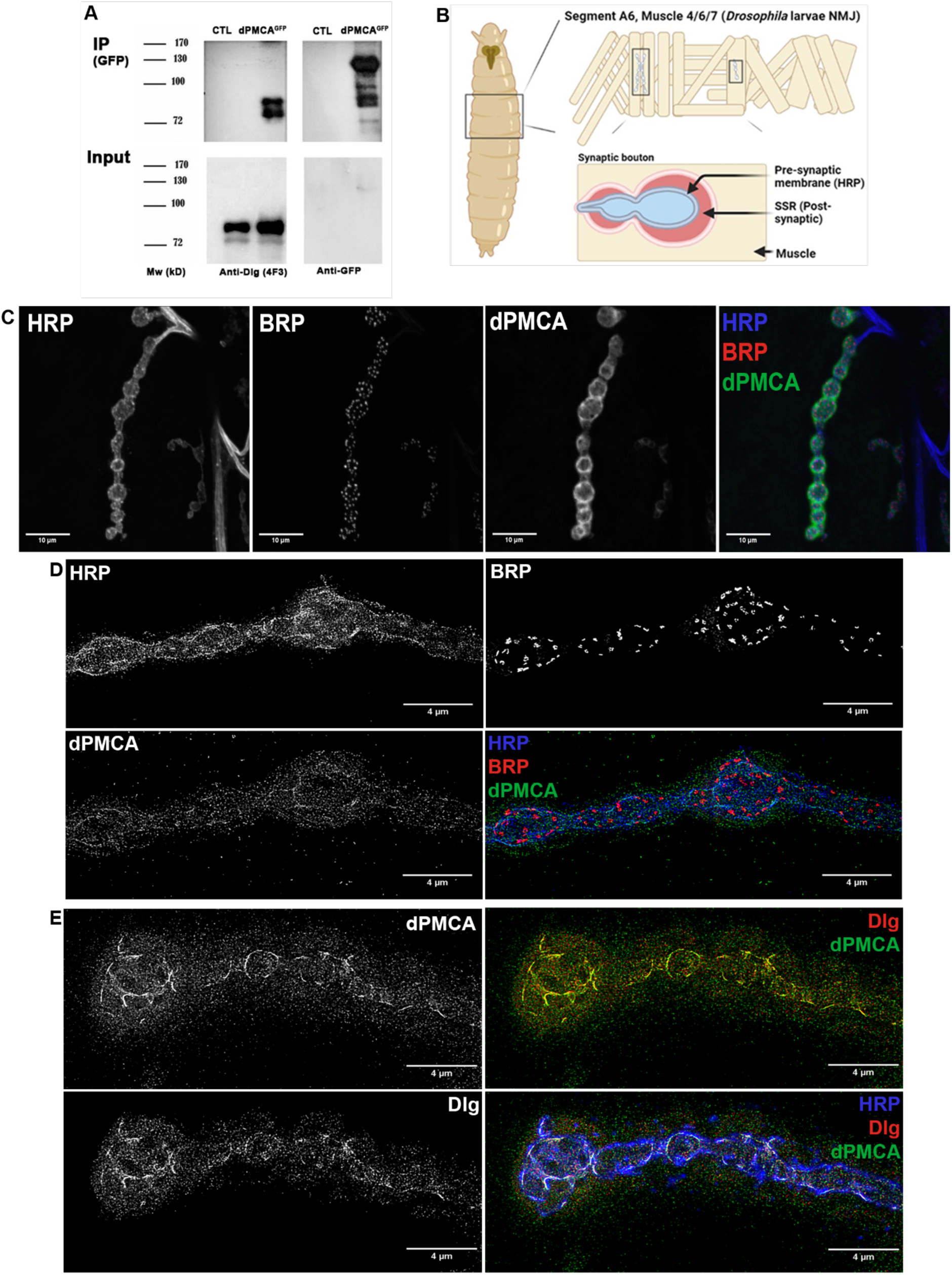
Dlg and PMCA associate and co-localize at presynaptic terminals of the *Drosophila* larval NMJ. **(A)** Co-immunoprecipitation (IP) of non-labelled endogenous Dlg with the genetically labelled PMCA-GFP from adult head extracts. Immunoprecipitation was performed using anti-GFP beads; immunoblots were probed with anti-Dlg (4F3) or anti-GFP antibodies. **B)** Schematic representation of the larval neuromuscular junction (NMJ), indicating the presynaptic bouton, active zones (AZs), and subsynaptic reticulum (SSR). **(C-D)** Confocal images (C) and (D) STED images of NMJs from animals expressing PMCA-RNAi and dlg-shRNA in the muscle cells to reduce postsynaptic signal and visualize the genetically labelled PMCA-GFP located in the presynaptic bouton. PMCA is observed to be enriched in membrane-associated patches that are largely excluded from Brp-positive (active zones) patches. **(E)** STED images of NMJs expressing endogenous Dlg-GFP and visualizing PMCA and HRP using labelled antibodies. Both proteins exhibit overlapping distributions within the presynaptic membrane. Pearson’s and Manders’ coefficients were calculated from the averaged presynaptic ROIs of 2 NMJ per larvae in 6 larvae. Pearson’s correlation index (mean ± SD) 0.801 ± 0.052; Manders co-distribution index: 0.972 ± 0.051; Manders co-distribution index (Threshold): 0.512 ± 0.136. Analysis restricted to neuronal membrane ROI (PMCA-Dlg): Pearson’s correlation index: 0.848 ± 0.02; Manders co-distribution index: 0.992 ± 0.01; Manders co-distribution index (Threshold): 0.471 ± 0.033. Scale bars: 4μm.

Previous studies have shown that PMCA is highly enriched in the subsynaptic reticulum (SSR), which surrounds presynaptic boutons, making it difficult to assess its presynaptic localization. To visualize the neuronal pool of PMCA, we reduced muscle PMCA-YFP expression using a muscle-specific GFP/YFP-RNAi approach on a *PMCA^YFP^/+* background and examined NMJs by confocal and STED microscopy. Under these conditions, PMCA-YFP was detected in discrete membrane-associated patches along the presynaptic bouton surface and was largely excluded from Brp-positive active zones (Fig. 1D; [16]).

To compare the distribution of PMCA-YFP and Dlg, we co-depleted both proteins in muscles and analyzed anti-GFP/anti-Dlg-labeled NMJs by STED microscopy. Both proteins showed highly overlapping distributions within presynaptic boutons (Fig. 1E). Quantitative colocalization analyses revealed high Pearson’s (0.801 +-0.052; n=6 larvae) and Manders’ coefficients (0.512 +-0,136; n=6 larvae), consistent with a close association between PMCA and Dlg at the presynaptic membrane. Together with the co-immunoprecipitation experiments, these observations indicate that Dlg and PMCA form part of the same molecular complex at presynaptic terminals

### Neuronal Dlg is required for normal presynaptic clustering of PMCA

To determine whether Dlg regulates PMCA localization, we analysed presynaptic endogenous PMCA-YFP expression in three *dlg* mutant backgrounds: *dlgS97*, *dlgA*, which specifically abolish DlgS97 or DlgA splice variants, respectively, and the hypomorphic *dlgXI-2* allele, which strongly diminishes both variants. Quantitative confocal analysis revealed a significant reduction of total NMJs PMCA fluorescence only in *dlgXI-2* mutant, whereas in *dlgS97* and *dlgA* mutants PMCA abundance was largely preserved (Fig. 2A-G)

**Figure 2:**
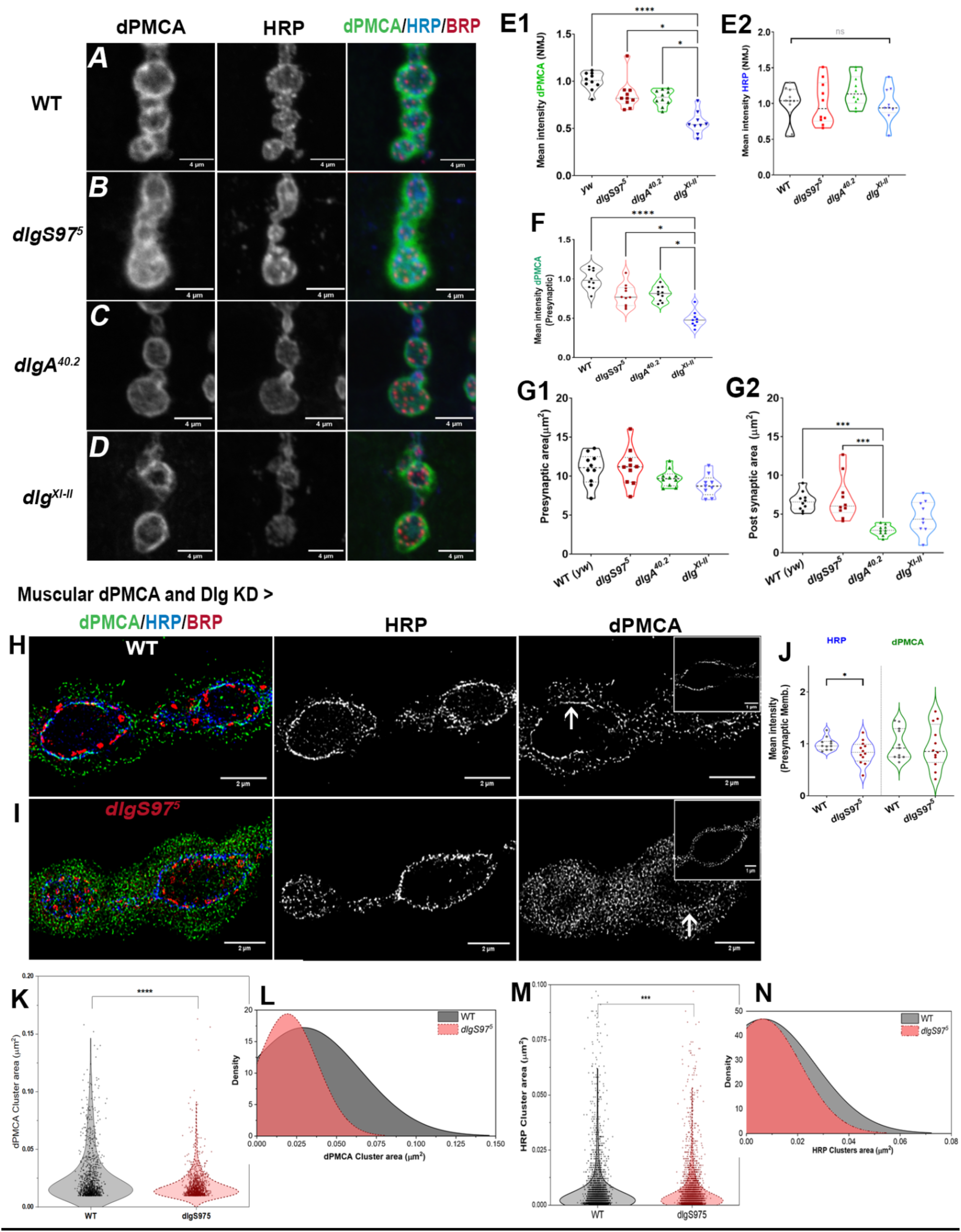
Neuronal Dlg is required for normal presynaptic clustering of PMCA. (A-D) Representative confocal images of NMJs from animals control, *dlgS97, dlgA*, and *dlgXI-2* mutants expressing PMCA-Venus and labelled using an anti-GFP (green) and anti-HRP (red). (E1-2) Quantification of the fluorescence intensity of PMCA and HRP for the whole NMJ across all mutants normalized to the control genotype. Only the double mutant shows a decrease in intensity of the PMCA labelling. (F) Comparison of the presynaptic PMCA, the *dlg* double mutant shows a decrease in the intensity of PMCA (G1-2), Quantification of the pre and post-synaptic areas occupied by PMCA. A decrease in the area occupied by PMCA in the postsynaptic in the double mutant could explain the lower PMCA fluorescence intensity observed. Kruskal-Wallis, Dunn, p-value: * < 0.05, ** < 0.01, *** < 0.001, **** < 0.0001. N = 9-11 larvae per condition (H-J) Representative STED images of presynaptic PMCA in control and dlgS97 mutant boutons after reduction of postsynaptic PMCA expression. <u>Inset in PMCA Channel</u>: Using HRP mask, the dPMCA signal in the presynaptic membrane was isolated - Note the fragmented cluster in *dlgS97* mutant (J) Quantification of synaptic PMCA and HRP levels indicate that in *dlgS97* mutant, overall PMCA abundance does not decrease in the presynaptic compartment. Mann-Whitney, p-value: * < 0.05. (K-N) Quantification of PMCA (K) and HRP (L) clusteŕs size and distribution in control and dlgS97 mutant larvae. Clusters <0.01 µm^2^ were discarded; clusters > 0.2 µm^2^ were added to the maximal range. The analysis shows that loss of neuronal DlgS97 alters the nanoscale organization of PMCA, reducing the size of the PMCA clusters without significant impact on its total presynaptic abundance. Mann-Whitney, p-value: *** < 0.001, **** < 0.0001.

Because the strong postsynaptic accumulation of PMCA masks presynaptic changes, we reduced postsynaptic PMCA expression and examined presynaptic PMCA organization using STED microscopy. Overall presynaptic PMCA-YFP fluorescence intensity or presynaptic area (assessed by HRP stain) did not differ significantly between control and *dlgA or dlgS97* mutant boutons, only in the hypomorphic mutant a clear decrease of the PMCA fluorescence intensity was observed. To assess PMCA distribution we set to analyse the synaptic boutons *of dlg* mutants by STED microscopy. Unfortunately dlgA and dlgXI-2 mutatn larvae were too sick to produce animals suitable for the technique. Therefore we performed the analysis with *dlgS97* mutants, the neuronal splice variant that produced larvae with less NMJ morphological defects. The analysis by STED microscopy, revealed clear alterations in PMCA organization. Specifically, while control boutons contained large membrane-associated PMCA clusters, *dlgS97* mutants displayed a more homogeneous distribution composed of smaller puncta (Fig. 2H-J).

Quantitative cluster analysis demonstrated a significant reduction in PMCA cluster size, along with an increase in the relative abundance of small clusters, in dlgS97 mutants (Fig. 2K-L), while the distribution of the Brp clusters remained separated from PMCA clusters. These findings indicate that neuronal DlgS97 is not required for trafficking presynaptic PMCA or for maintaining its levels but it is necessary for the assembly and/ or maintenance of PMCA clusters within peri-active-zone membrane domains.

On the other hand, at the postsynaptic side, the area of the postsynaptic NMJ was reduced in *dlgA* mutants (as previously shown[14], [21], [22], [23]) (Fig. 2G). Consistently; muscle-specific knockdown of Dlg (shRNA-*dlg*) produced a significant decrease in NMJ PMCA staining, supporting the association of Dlg-PMCA postsynaptically as well as presynaptically (Supp. Fig. 2E–L).

### Disruption of Dlg-dependent PMCA organization alters calcium-dependent neurotransmitter release

Given the established role of PMCA in calcium regulation at the larval NMJ, we next examined whether disruption of Dlg-dependent PMCA organization correlates with the changes in synaptic calcium dynamic. To test it, we recorded the evoked and spontaneous synaptic currents by two-electrode voltage clamp in dlg mutants and in animals expressing neuronal or motoneuronal RNAi against *dlg* or *pmca* (Fig. 3A).

**Figure 3.**
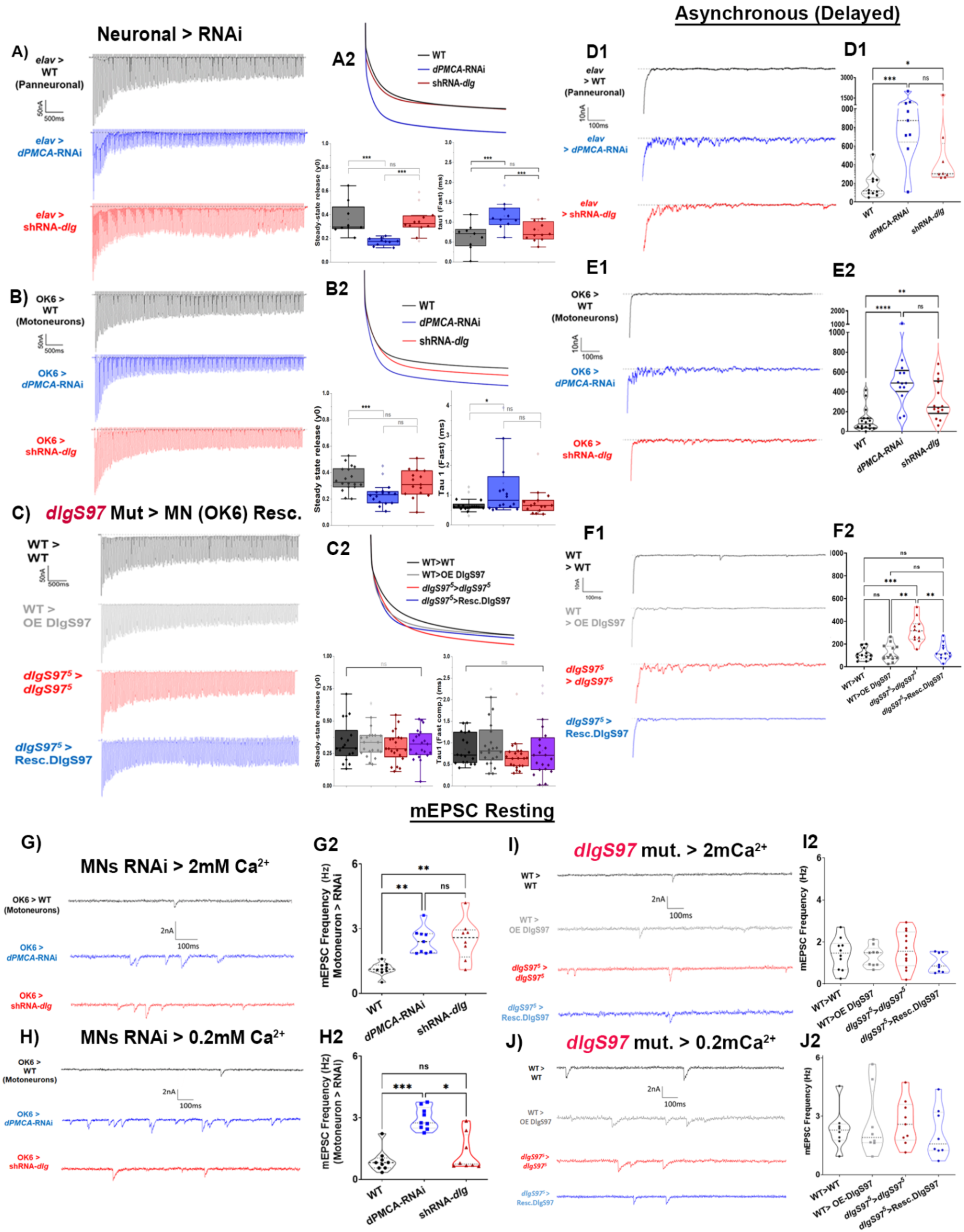
Dlg and PMCA regulate calcium-dependent neurotransmitter release. (A-C) Representative EPSC trains recorded during 150 stimuli at 20 Hz in control (*elav-Gal4/+*) and panneuronal *expression* (ElaV) (A) or motoneuronal expression (*OK6*) (B) of PMCA-or dlg-specific RNAi effectors; (C) the *dlgS97* mutant (*dlgS97^5^*) with their respective controls and motoneuronal (OK6) conditions are also presented. The amplitude of each EPSC was normalized to the amplitude of the first EPSC to generate a graphical representation of each recording which was then fitted to a 2-exponential decay function to generate the respective curves for each condition represented in A2 and B2 maintaining the same colour code as in A and B (gray: control; blue: down regulation of PMCA in neurons; and red: down regulation of *dlg* in neurons). (A2-C2) The average and dispersion of the parameter y0 (the amplitude of the EPSC at steady state) from each fitted curve of all the recordings obtained and the average and dispersion of the parameter tau1 (the time constant of the fast component) from each fitted curve of all the recordings obtained for each genotype. (D-F) Representative recordings of the asynchronous release after the tetanic stimulation for panneuronal (D) and motoneuronal (E) knockdown fo*r dlg* and *pmca* and for the *dlgS97* mutant background conditions (F). On the right of each recording is shown the quantification of the total charge transferred (in nC) in 300 ms, 30 ms after the end of the stimulus. (G-J) Representative recordings of spontaneous synaptic currents (mEPSC) and the quantification of its frequency under low (0.2 mM) (H,J) and physiological (2 mM) (I,J) HL3.1 extracellular calcium conditions. Neuronal reduction of either PMCA or Dlg increased the frequency of the mEPSCs at 2 mM calcium concentration, consistent with altered presynaptic calcium regulation. However, under less stringent conditions, only PMCA downregulation showed an altered mEPSC frequency. Data points represent the average obtained in one larva. Kruskal-Wallis, Dunn, p-value: * < 0.05, ** < 0.01, *** < 0.001, **** < 0.0001. N = panneuronal: 7-11, Motoneuron: 9-11, and dlgS97 mutant: 12.

At physiological extracellular calcium (2 mM) and during repetitive stimulation (150 stimuli at 20 Hz), cytosolic calcium accumulates generating a rapid decrease of the amplitude of the evoked postsynaptic currents (EPSC). This decrease is known as depression and its kinetics can be fitted to an exponential equation with two or more parameters. The depression curves were fitted with at least two exponentials for which we compared the tau of the fast component for all genotypes as well as the amplitude at the steady state. Neuronal downregulation of PMCA displayed a pronounced increase of EPSC depression relative to controls, reflected as a smaller amplitude at steady state, although with a slower tau (Fig. 3A-B, Blue). However, following neuronal reduction of Dlg and in dlgS97 mutants (Fig. 3A-C, Red) no differences on either parameter was observed. These results indicate that PMCA is needed to maintain the calcium homeostasis during a repetitive stimulus.

Once tetanic stimulation ceases, the accumulated calcium results in neurotransmitter release that is highly asynchronic and lasts until the calcium returns to basal. Therefore, we determined the residual presynaptic calcium after repetitive stimulation, quantifying the asynchronous release in control animals and with neuronal PMCA knockdown. We observed a marked increase in asynchronous neurotransmitter release in PMCA neuronal knockdown compared with controls (Fig. 3D-E). A significant increase was also observed in dlgS97 mutants and in animals with neuronal Dlg knockdown. Expression of DlgS97 in motoneurons rescued the mutant phenotype (Fig.3F), supporting a presynaptic role for DlgS97 in this process. These findings are consistent with altered regulation of activity-dependent calcium signals when PMCA function or organization is disrupted.

We next examined spontaneous neurotransmitter release. Changes in extracellular calcium modify the frequency of the spontaneous neurotransmitter release due to changes in the minimal concentration of calcium that the synapse can maintain. Under physiological extracellular calcium conditions (2 mM), neuronal reduction of either PMCA or Dlg significantly increased mEPSC frequency (Fig. 3I-J). However, at low extracellular calcium (0.2 mM) (Fig.3G-H), PMCA knockdown continued to elevate spontaneous release frequency, whereas Dlg reduction had little effect. Notably, *dlgS97* mutant did not replicate the phenotype observed with the presynaptic downregulation of *dlg*, suggesting compensatory mechanisms or the requirement of dlgA for this function. Together, these observations support an essential role for PMCA in the control of resting calcium (and by that synaptic activity) and support a Dlg-dependent-PMCA clustering that makes the function of PMCA more efficient, but it is not absolutely required for its function.

### PMCA contributes to EPSC decay kinetics during sustained stimulation

In vertebrate synapses, PMCA activity is generally associated with bulk calcium clearance. Recent work, however, has uncovered an unexpectedly fast pump activity of vertebrate PMCA ([24] [25]), but its activity has been associated mostly with bulk calcium clearance rather than rapid regulation of active-zone calcium. To assess whether PMCA influences fast components of synaptic transmission at the *Drosophila* NMJ, we analysed individual EPSC decay kinetics along the repetitive stimulation.

Across all genotypes, EPSC decay times became progressively faster during the initial phase of the stimulus train and subsequently reached a stable plateau (Fig.4A-C). However, neuronal PMCA knockdown consistently prolonged EPSC decay kinetics relative to controls throughout the stimulation train (Fig. 4A-B), supporting a slower clearance of the fast active-zone calcium. Significant effects were observed for the first, the fifth, and the seventy-fifth EPSCs (Fig.4D-G). In contrast, neither neuronal Dlg reduction nor the dlgS97 mutation produced detectable changes in EPSC decay kinetics (Fig. 4D-G, H-I).

**Figure 4.**
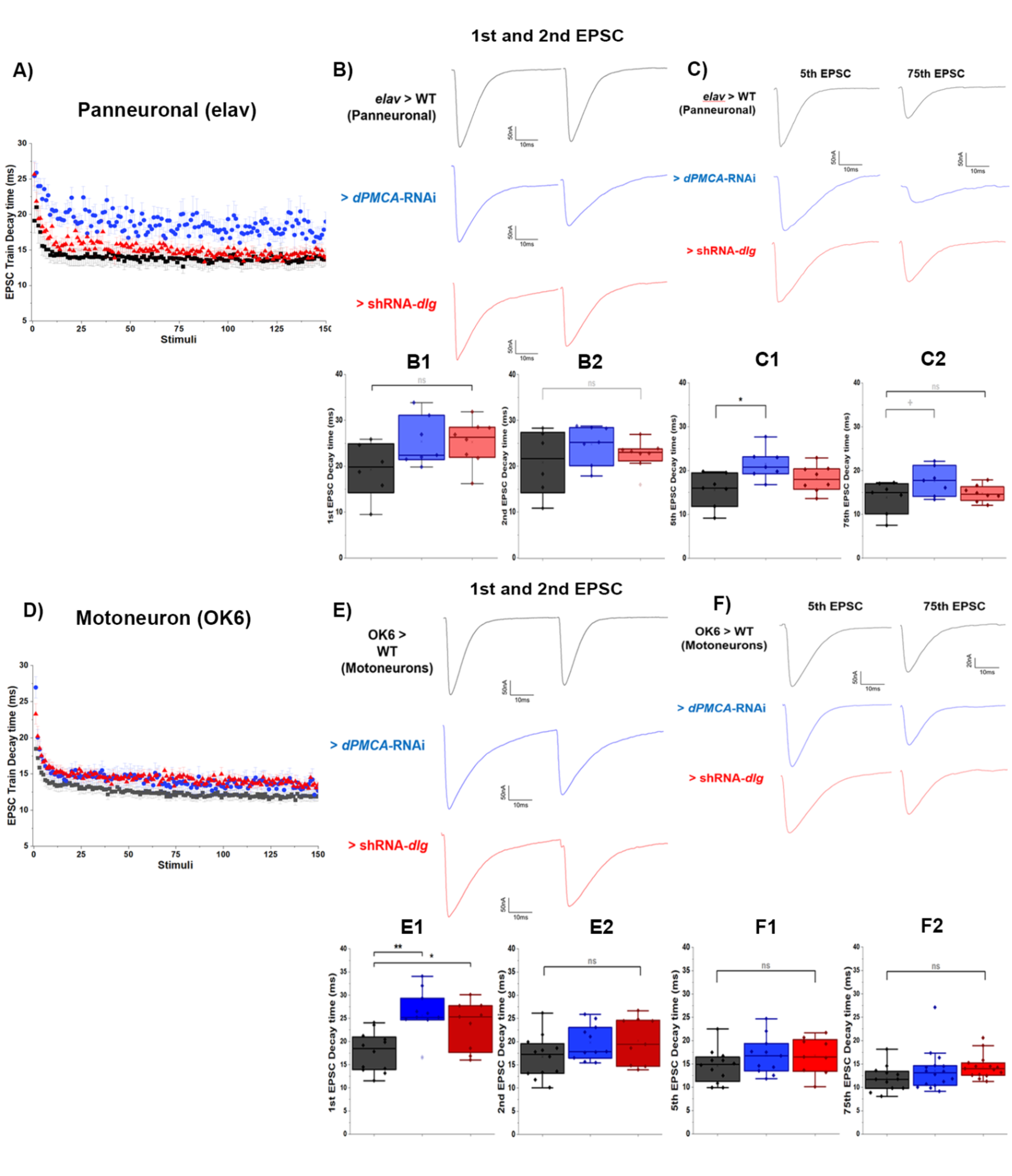
PMCA contributes to the evoked postsynaptic current (EPSC) decay kinetic during the repetitive stimulation. **(A,D)** Temporal change of the EPSC decay times throughout the train of stimulus in control animals (black dots), neuronal downregulation of PMCA (blue dots) and neuronal downregulation of *dlg* (red dots). **(B,C) (E,F)** Representative EPSCs recorded at selected positions within the stimulus train (1st, 2nd, 5th, and 75th stimuli) for panneuronal **(B,C)** and Motoneuronal **(E,F)**. The corresponding decay-time quantifications for each representative recording are presented in the boxplots (B,C1-2) (E,F1-2). Reduction of PMCA prolonged EPSC decay kinetics, whereas reduction of Dlg had little effect on this parameter. The data represent individual larvae. Statistical analyses are described in Methods. Kruskal-Wallis, PostHoc Dunn, p-value: + = 0.1, * < 0.05, ** < 0.01. Larvae per condition = panneuronal: 7-11, Motoneuron: 9-11, and dlgS97 mutant: 12.

These findings indicate that PMCA contributes to mechanisms that shape the temporal profile of synaptic transmission during repetitive activity by controlling fast and high affinity calcium concentration in the synapse and that the clustering of PMCA is not necessary for this function.

### PDZ-mediated interaction with Dlg promotes presynaptic clustering of PMCA

To investigate the molecular basis of the Dlg-PMCA interaction, we analysed reporter constructs containing PMCA endogenous alternative C-terminal tails. These reporters differed in the presence or absence of canonical PDZ-binding motifs (Fig. 5A). Co-immunoprecipitation experiments demonstrated that reporters containing either canonical or non-canonical PDZ-binding motifs associated with endogenous Dlg, whereas the reporter lacking a PDZ-binding motif did not (Fig. 5B).

**Figure 5.**
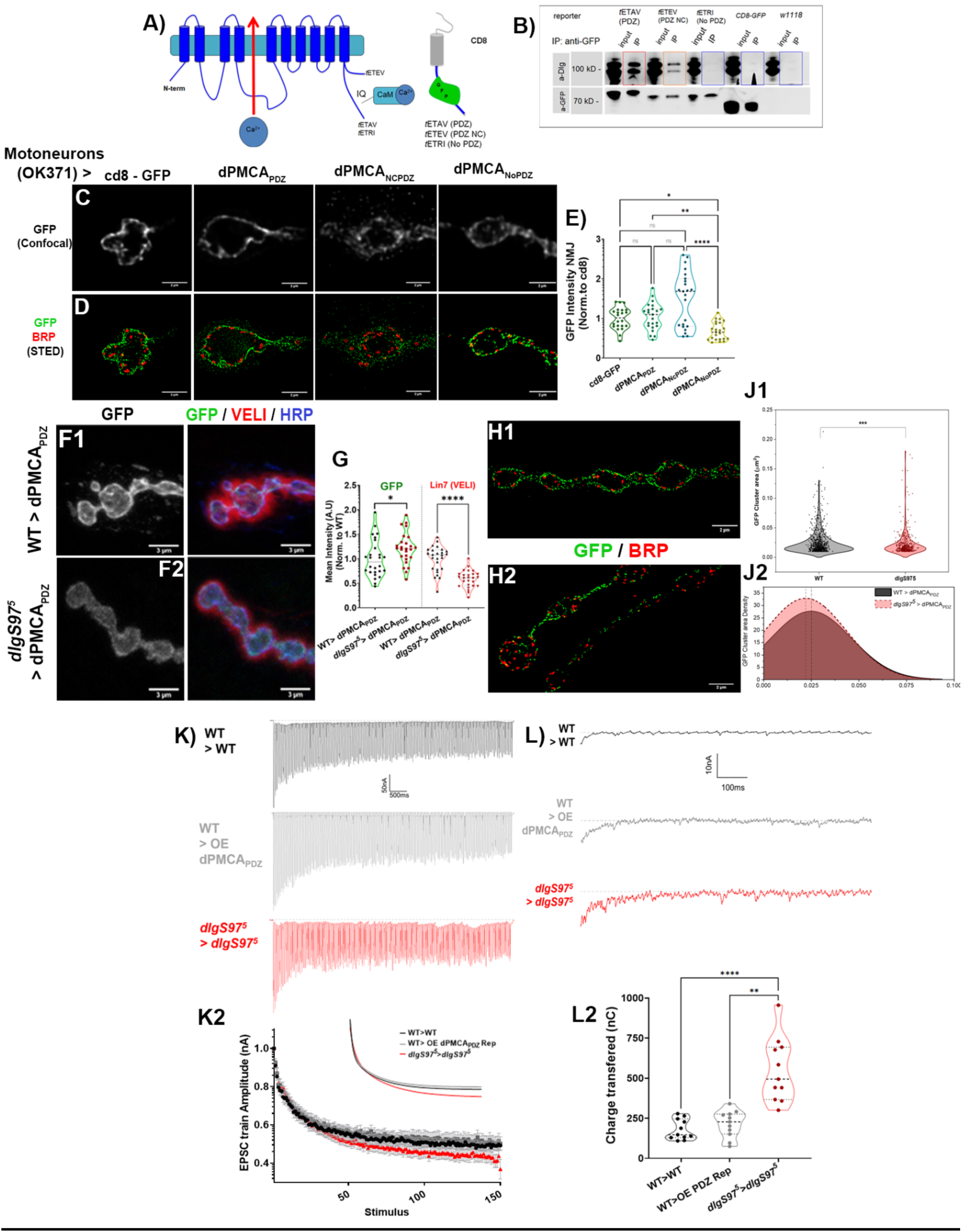
PDZ-mediated interaction with Dlg promotes clustering of PMCA C-terminal reporters. **(A-B)** Schematic representation of PMCA C-terminal reporter constructs containing PDZ-binding (PDZ), non-canonical PDZ-binding (NcPDZ), or non-PDZ-binding (NoPDZ) motifs. **(C)** Co-immunoprecipitation of reporter proteins with endogenous Dlg from Drosophila Heads. **(D-E)** Confocal and STED images of NMJ expressing PMCA C-terminal tail reporters in motoneurons. PDZ-containing reporters display clustered presynaptic distributions, whereas the NoPDZ reporter is distributed more uniformly. Quantification of reporter expression levels was normalized to the control reporter (CD8-GFP) for comparison. **(F-G)** Representative confocal images of NMJ synaptic terminals immunostainings of control and *dlgS97* mutant larvae showing the reporter (green), the Dlg-associated protein Lin7 (VELI, red), and the presynaptic membrane marker HRP. **(G)** Quantification of average fluorescence intensity in samples expressing PMCA-PDZ reporter. The fluorescence of the reporter and of the protein Lin7 were normalized to the control (WT). Kruskal-Wallis, PostHoc Dunn, p-value: * < 0.05; ** < 0.01, ****<0.0001; n= 24 NMJ (12 larvae). **(H-J) (H)** Representative STED images of NMJ synaptic terminals immunostained of control and *dlgS97* mutant larvae showing the reporter (green) PMCA-PDZ and the active zone protein Bruchpilot (red). **(J)** Quantification of the size (J1) and density (J2) of the presynaptic PDZ-reporter clusters. Mann-Whitney, p-value: *** < 0.001. **(K-L)** (K) Electrophysiological EPSC recordings of PDZ reporter expression in control (Black), Overexpression (OE) (Gray), and dlgS97 mutant (Red) during repetitive stimulation. (K1) For each condition, every train was normalized to the first EPSC amplitude and adjusted to a fitting function (2-exponential decay) to generate the respective curves for each condition. The adjusted fitting curve parameters steady-state release (y0), Amplitude - slow component (A2), Tau1, and Tau2 (Fast and slow decay component - not presented) obtained from each train were calculated (O) Kruskal-Wallis, PostHoc Dunn, p-value: ns; n= 24 NMJ (12 larvae). (L) Representative recordings and quantification (L2) of asynchronous (delayed) release (in nC) of Control, OE, and *dlgS97* mutant after tetanic stimulation. Kruskal-Wallis, PostHoc Dunn, p-value: ****<0.0001; n= 24 NMJ (12 larvae).

When expressed in motoneurons, the PDZ-containing reporter displayed a clustered distribution that closely resembled endogenous PMCA (Fig.5C-E) - In contrast, the non-PDZ reporter exhibited a more continuous and uniform membrane distribution as well as a lower fluorescence intensity (Fig.5D).

To determine whether Dlg was required for reporter clustering, we examined the PDZ reporter in a *dlgS97* background. Agreeing with our previous results, loss of DlgS97 did not decrease the fluorescence of the reporter in the button (Fig 5E), however, it significantly reduced reporter clustering and shifted the distribution toward smaller puncta (Fig. 5F-L). Thus, PDZ-mediated interaction with Dlg is required for normal assembly of PMCA-PDZ containing membrane clusters. As an internal control, mutant NMJs were labeled with anti-Veli antibodies. Veli forms a complex with DlgS97 that is highly dependent on the presence of DlgS97, therefore, in the dlgs97 mutant the fluorescence intensity of Veli is highly diminished [22].

Finally, overexpression of the PDZ reporter did not interfere with PMCA function, assessed by the asynchronous release and the depression constants during repetitive stimulation in WT background. These results indicate that the PDZ motif could not compete with the endogenous PMCA proteins, suggesting that the clusters might form in a structure where the PDZ binding site becomes not accessible for recruiting new molecules.

## DISCUSSION

Calcium extrusion is a critical determinant of presynaptic calcium homeostasis, yet the molecular mechanisms controlling the calcium extrusion remain not completely understood. In this study, we determine that the MAGUK scaffold protein Dlg is necessary for the presynaptic clustering of PMCA at the *Drosophila* larval neuromuscular junction. We show that Dlg and PMCA are part of the same molecular complex *in vivo*, that neuronal Dlg is required for the formation of normal presynaptic PMCA clusters, and that disruption of this organization is associated with altered calcium-dependent neurotransmitter release. Together, these findings identify a previously unrecognized role for Dlg in the organization of presynaptic calcium extrusion mechanisms.

The interaction between PMCA proteins and Dlg family members has previously been reported in mammalian non neural culture cells, where PDZ-mediated interactions contribute to PMCA localization and membrane clustering [17], [31]. However, evidence linking this interaction to synaptic function *in vivo* has remained limited. Our co-immunoprecipitation and STED microscopy analyses demonstrate that Dlg and PMCA are closely associated within presynaptic boutons of the *Drosophila* NMJ. Importantly, loss of DlgS97 did not substantially reduce presynaptic PMCA abundance but instead altered its nanoscale organization, converting large membrane-associated clusters into smaller and more uniformly distributed structures (Fig.2). These observations indicate that neuronal Dlg primarily regulates the spatial arrangement of PMCA rather than its traffic to the synaptic terminals.

The functional significance of this organization became evident from our electrophysiological analyses (Fig.3-4). Reduction of either PMCA or neuronal Dlg increased spontaneous neurotransmitter release under physiological resting conditions and enhanced asynchronous release following tetanic stimulation. Because both parameters are highly sensitive to residual presynaptic calcium, these results are consistent with altered calcium homeostasis in terminals lacking either PMCA function or proper PMCA organization. Notably, the phenotypes produced by Dlg reduction were consistently milder than those observed following PMCA knockdown. This distinction suggests that Dlg is not required for PMCA activity per se, but rather enhances its effectiveness through spatial organization within the presynaptic membrane, although we cannot discard that the physical interaction per se might result in an improved efficiency of the pump. We therefore propose that Dlg-dependent clustering optimizes the ability of PMCA to regulate physiologically relevant calcium signals generated during synaptic transmission.

PMCA proteins are generally regarded as high-affinity transporters that contribute to the restoration of resting intracellular calcium [27], [32], [33], [34]. Our results support this established function, as a reduction in neuronal PMCA increased spontaneous release frequency even under low extracellular calcium conditions. Additional support for the function of PMCA in the regulation of resting calcium is the result showing that Dlg and PMCA knockdown increased post-tetanic asynchronous release and accelerated synaptic depression. These findings are consistent with previous work identifying PMCA as a major contributor to bulk calcium clearance at the *Drosophila* NMJ [13] , [35] and further suggest that its spatial organization is an important determinant of calcium-dependent synaptic function.

Interestingly, neuronal knockdown of PMCA additionally slowed the decay kinetics of the EPSC, whereas reduction of Dlg had little effect on this parameter. This result suggests a role of PMCA in the fast calcium dynamic at the active zone. Although our study did not directly measure presynaptic calcium, electrophysiological synaptic parameters have been shown to follow active zone calcium. Moreover, despite additional mechanisms, including local calcium buffering, endoplasmic reticulum sequestration, mitochondrial uptake, or other calcium transport systems [36] , [37] , [38] , [39] , [40] , [41] (ref), may also contribute to shaping release kinetics, the decrease in PMCA levels associated to its genetic manipulation is the more plausible mechanism to interpret the results.

The analysis of PMCA C-terminal reporters provides a mechanistic explanation for the organization of PMCA clusters. Reporters containing PDZ-binding motifs associated with endogenous Dlg and recapitulated the clustered distribution of presynaptic PMCA, whereas a reporter lacking a functional PDZ-binding sequence failed to do so. Furthermore, the clustered organization of the PDZ reporter was disrupted in *dlgS97* mutants, closely paralleling the phenotype observed for endogenous PMCA. These observations support a model in which PDZ-mediated interactions recruit PMCA into Dlg-associated scaffold complexes within peri-active-zone membrane domains. The concomitant reduction in Lin-7/VELI levels observed in *dlgS97* mutants (Fig.5F-H), further suggests that PMCA may be incorporated into larger scaffold assemblies that organize presynaptic membrane specializations.

At the synaptic terminal, several different mechanisms act in concert to maintain calcium homeostasis. The mammalian PMCA has been described as a high-affinity, low-efficiency system with a secondary role in the removal of Ca^2+^ in microdomains [13] , [26] , [27] , [28] [29] , [30] , leaving the role of calcium extraction near the active zone and during activity to a faster transport mechanism that remains to be characterized. However, it is known that response speed and efficiency depend on the PMCA isoform [42] and on binding to modulatory proteins such as neuroplastin. PMCA associated with neuroplastin reaches rates comparable to those of the Na^+^/Ca^2+^ exchanger (5000 cyc/s; [43]). In *Drosophila*, the affinity and transport rate of the pump are unknown; interpretations of its function are based on available data from mammalian PMCA. Here, we have explored the involvement of the PMCA in the regulation of the presynaptic calcium and its association with the main scaffold in the *Drosophila* NMJ.

Our findings support a model in which Dlg acts as a presynaptic scaffold, organizing PMCA into specialized membrane domains that support resting calcium homeostasis and thereby regulate calcium-dependent neurotransmitter release. Disruption of this organization redistributes PMCA within the membrane, increasing spontaneous and asynchronous neurotransmitter release. Furthermore, we demonstrate that PMCA contributes in vivo to the rapid restoration of calcium levels following a single stimulus, a process previously considered too fast to be mediated by the pump, in agreement with recent measurements of PMCA kinetics [43]. Because this function does not require Dlg or PMCA clustering, it may involve an alternative membrane-anchoring mechanism that warrants further investigation. Overall, our findings highlight the nanoscale organization of calcium extrusion machinery as an important regulator of synaptic physiology and establish the Dlg–PMCA complex, together with PMCA itself, as a key determinant of presynaptic calcium homeostasis at the Drosophila neuromuscular junction, both at rest and during neuronal activity.

## Materials and Methods

### Fly strains and maintenance

the yw strain was used as a control for *dlg* mutants; each GAL4 line crossed to the *w1118* line was used as a control for all other experimental lines. flies. PMCA mutant and interference RNA for PMCA and Dlg, as all Gal4 line sources are detailed in Table 1 in the annexes. Flies were maintained and crossed at 25°C; for the induction of Gal4, UAS-carrying dsRNA were transferred to 29°C during the third instar.

### Immunoprecipitation assays

For the CoIP immunoprecipitation experiments, *Drosophila melanogaster* adult heads were used as described [32], heads from 5 mL of flies frozen with liquid nitrogen were separated from the bodies by vortexing the tube at maximum speed, and then the flies were passed through a sieve. The recovered heads were homogenized in 500 µL of 1x NP-40 buffer in the presence of protease inhibitor cocktail. The homogenized was centrifuged at 14,000 rpm, and the supernatants were used for the IP. Proteins were quantified by the Bicinchoninic acid method. 10% of the sample was saved and stored at - 20°C for use as a loading control. 25 µL of Magnetic Beads (per 2 mg of protein) coated with anti-GFP (GFP-trap magnetic agarose, ChromoTek) were added to the rest of the sample, diluted with 175 µL of 1X NP-40 buffer, and incubated for 4 to 6 hours. The beads were magnetically separated, washed, and eluted with 30 µL of 1x loading buffer, incubated for 5 minutes, and then heated at 95°C for 5 min. The samples were loaded in a 10% acrylamide SDS gel, transferred to nitrocellulose, and revealed with anti-Dlg or anti-GFP.

### Immunohistochemistry, super-resolution (STED), and confocal fluorescence microscopy

**Male third-instar** larvae and 5-7-day-old adult brains were used [50], [51], [52]. Primary anti-GFP (Rabbit, Invitrogen; 1:100), anti-Bruchpilot (Brp - Nc82; 1:300, Mouse, DSHB), PMCA (Rabbit, G. Lnenicka 56; Rabbit, Ulrich Thomas, LIN), anti-RFP, and membrane Fasciclin II ( DSHB 1:100). For STED microscopy, Atto-580 and Alexa 647N-conjugated anti-GFP and anti-RFP nanobodies (Invitrogen), were used. Secondary antibodies conjugated with the DyLight Alexa fluorophores 405, Cy3, Rhodamine, FITC, Cy5, and Anti-HRP (1:200) (Donkey, Alexa fluor 405 and Alexa fluor 647N - Jackson laboratories) were used to obtain a fluorescent signal. Samples were mounted in Vectashield medium (Vector Laboratories, Burlingame, CA).

Confocal microscopy image stacks (optical sections 0.2 - 0.5 µm) of NMJs on muscles 4, 6, and 7 in segments A2 or A3 and the ventral nerve cord of dissected larvae were obtained on Olympus FV1000, Olympus BX61WI, and Leica SP8 Lightning microscopes, 60x objective, N.A. 1.4. For adult brains, whole-brain images and MB calyces were obtained on an Olympus FV1000 using 60x magnification and optical sections of 0.3 to 0.5 μm. STED super-resolution microscopy images were performed at the Leibniz Institute for Neurobiologie (LIN) using a Leica SP8-3X microscope with a 93x objective and 0.13 μm sections, using 488, 561, and 633 nm lasers; the depletion lasers used were 660 and 770 nm. Subsequently, the images were analyzed using the Fiji software (ImageJ) version 1.51. Deconvolution of STED and confocal microscopy images was performed in Huygens software; parameters for confocal: Maximum iterations 60, Signal / Noise: 30, quality iterations 0.01; STED parameters: maximum iterations 60, signal/noise 15, quality iterations: 0.001. For the design of three-dimensional images, the Imaris software (standard configuration) was used.

### Analysis of fluorescence intensity, colocalization, and cluster area

The deconvolved images labelled with anti-HRP were used to outline the presynaptic compartment. anti-BRP was used to label the active zones and C57-GAL4> mCherry-CAAX to delimit the postsynaptic SSR. For adult brains, MB Calyxes ROIs were generated using the combination of DAPI, PMCA-Cherry, BRP, or dAlphaNAchR masks.

The Fiji/ImageJ “Threshold-Otsu” feature was used to generate masks containing the fluorescence associated with anti-HRP. The regions of interest (ROIs) were generated for each bouton and used to quantify the fluorescence intensity in the images; the data were collected in Microsoft Excel for subsequent statistical analysis. For the colocalization tests, we used the JaCoP function (Fiji/ImageJ), restricting the analysis to the presynaptic ROI or to ROIs built from other labels, and excluding signals outside the ROI. For each comparison between 2 channels, the Pearson correlation (r) and Manders co-distribution (M1, M2, m1, m2) were obtained. For the cluster area analysis, the bouton was outlined using the HRP signal and restricted to 200 nm from the outer boundary. With this outline we generated a mask that limited the analysis to the neuronal membrane for each Z-optical slice. The mask for each slice was compared to the original confocal and STED image. In deconvolved STED microscopy images we used the neuronal membrane mask and inside that space generated ROIs for each particle. For each particle we obtained the area. Then an histogram was generated considering the size of the particles. The analysis was restricted to particles larger than 0.01 μm^2.

### Electrophysiology

Postsynaptic currents of third instar larvae were recorded in segment A3 of ventral longitudinal muscle 6, using the two-electrode Voltage Clamp technique (OC-725C, Harvard Apparatus, Holliston, MA). The larvae and electrode pipettes will be immersed in the HL3.1 medium (in mM: 10 NaHCO3, 5 KCl, 4 MgCl, 5 HEPES, 70 NaCl, 5 Trehalose, 115 Sucrose, pH 7.4), in low and high calcium (0.2 mM and 2 mM, respectively). For the asynchronous release analysis, recordings were made for 100 seconds before and 100 seconds after the high-frequency train (150 stimuli at 20Hz). Using a stimulator (Master 8, A.M.P.I., Jerusalem, Israel), the nerve innervating the recorded muscles was stimulated with the minimal current necessary to obtain a response in the postsynaptic currents.. Acquisition was performed using WinEDR software and analysis of electrophysiological responses using Clampfit 9.0 (Molecular Devices, Sunnyvale, CA).

For the analysis of the kinetics of evoked currents (EPSCs) during the neuronal stimulation train, the maximum amplitude and the decay time were quantified using the Clampfit 9.0 program and the event detection function the kinetics of the stimulation trains were analyzed for each experimental condition, adjusting to an exponential of 2 curves (y = A1 * exp (-x / t1) + A2 * exp (-x / t2) + y0) and then the Tau’s of each curve were obtained. The graphs show t1 and y0.

To quantify the asynchronous release after the high-frequency train, we obtained the total postsynaptic current in nanocoulombs (nC) from 30 ms after the start of the last stimulus in the stimulation train to 300 ms, quantifying the area under the baseline during the period. The analysis of the frequency and amplitude of spontaneous events was carried out in Mini-analysis (Synaptosoft); the number of events and the amplitude of each were analyzed over 10 minutes of recording.

### Statistical analysis

Each data set corresponded to the average quantification of 3 to 6 boutons obtained from two images of NMJ branches per larvae (NMJ muscle 4/6/7 left and right side, respectively). The number of data corresponds to individual larvae (2 images per larva, 3-6 boutons per image, analyzed per larva). Mann-Whitney (non-parametric) test was used for control analysis versus the RNAi/mutant condition, and Kruskal-Wallis ANOVA (non-parametric), with Dunn Post Hoc test, to compare control versus different RNAi/mutant conditions.

---

**Annex Table 1:** Fly strains used in this investigation.

| **Mutants/Deficiency strains** |  |  |
|---|---|---|
| *y*,*w*, *dlgA<sup>40.2</sup>*/FM7 | Sierralta Lab | X |
| *y*,*w*, *dlg<sup>XI/2</sup>* /FM7 | Sierralta Lab | X |
| *y*,*w*, *dlgS97<sup>5</sup>* | Sierralta Lab | X |
| PMCA-MiMIC12515/ In (4) (60856) | Bloomington | IV |
| **GFP/RFP-Trap** |  |  |
| Dlg-GFP (115-015 A133-620) | Kyoto | X |
| PMCA-GFP (P(Element) on PMCA (B215-109)) | Kyoto | IV |
| PMCA-mCherry (PMCA -MiMIC-mCherry) | BestGene / Sierralta Lab | IV |
| **Gal4-UAS Strains** |  |  |
| *elav*-GAL4 (Pannneuronal) |  |  |
| yw, *dlgS97<sup>5</sup>* , *elav*-GAL4 | Sierralta Lab | X |
| yw, *dlgA*<sup>40.2</sup>, *elav*-GAL4 | Sierralta Lab | X |
| OK6-GAL4 | Sierralta Lab | X |
| yw, *dlgS97<sup>5</sup>*; OK6-GAL4 | Sierralta Lab | II |
| yw, *dlgA*<sup>40.2</sup>; OK6-Gal4 | Sierralta Lab | X, II |
| yw, *dlg<sup>XI/2</sup>*; OK6-GAL4 | Sierralta Lab | X, II |
| *w<sup>1118</sup>* | Sierralta Lab | X, II |
| CantonS | Sierralta Lab | X |
| *y<sup>1</sup>w<sup>1</sup>* (yw) | Sierralta Lab | - |
| UAS-DlgS97--GFP | Sierralta Lab | II |
| UAS-DlgA-GFP | Sierralta Lab | II |
| UAS-RNAi-TRiP-Dlg1 /TM3 (33620) | Bloomington | III |
| UAS-RNAi-TRIP-Dlg1 /TM3 (34854) | Bloomington | III |
| UAS-RNAi-TRiP-PMCA (31572) | Bloomington | III |
| UAS-RNAi-PMCA (VDRC: 30203GD) | VDRC | III |
| UAS-RNAi-PMCA (VDRC: 101743) | VDRC | II |
| UAS-RNAi-PMCA (VDRC: 109188) | VDRC | II |
| UAS-CTerm-PMCAPDZ-cd8GFP (E10) /CyO | Thomas Lab | II |
| UAS-CTerm-PMCANcPDZ-cd8GFP (D4) /CyO | Thomas Lab | II |
| UAS-CTerm-PMCANoPDZ-cd8GFP (F1) /TM6b | Thomas Lab | III |

## Notes

### Competing Interest Statement

The authors have declared no competing interest.

### Funding

- Fondecyt 1210586 (Chile)

